# Genetic analyses reveal population structure and recent decline in leopards (*Panthera pardus fusca*) across Indian subcontinent

**DOI:** 10.1101/746081

**Authors:** Supriya Bhatt, Suvankar Biswas, Krithi K. Karanth, Bivash Pandav, Samrat Mondol

## Abstract

Large carnivores maintain the stability and functioning of ecosystems. Currently, many carnivore species face declining population sizes due to natural and anthropogenic pressures. The leopard, *Panthera pardus*, is probably the most widely distributed and adaptable large carnivore, still persisting in most of its’ historic range. However, we lack subspecies level data on country or regional scale on population trends, as ecological monitoring approaches are difficult to apply on such wide-ranging species. We used genetic data from leopards sampled across the Indian subcontinent to investigate population structure and patterns of demographic decline. Our genetic analyses revealed four distinct subpopulations corresponding to Western Ghats, Deccan Plateau-Semi Arid, Shivalik and Terai region of north Indian landscapes, each with high genetic variation. Coalescent simulations with 13 microsatellite loci revealed a 75-90% population decline in between 120-200 years ago across India, possibly human induced. Population-specific estimates of genetic decline are in concordance with ecological estimates of local extinction probabilities in four sub-populations obtained from occupancy modelling of historic and current distribution of leopards in India. Our results confirm population decline of a widely distributed, adaptable large carnivore. We re-iterate the relevance of indirect genetic methods for such species, and recommend that detailed, landscape-level ecological studies on leopard populations are critical to future conservation efforts. Our approaches and inference are relevant to other widely distributed, seemingly unaffected carnivores such as the leopard.

## Introduction

Large carnivores are critical to ecosystem structure and functioning (Sergio et al. 2008) and their absence can lead to significant changes in trophic cascades (Terborgh et al. 2001, Steneck 2005, Estes et al. 2011, Ripple et al. 2014). Growing natural and anthropogenic pressures in the form of climate change, habitat and prey depletion, poaching and human-wildlife conflicts are pushing large carnivores into ever-shrinking habitat islands and severely exacerbating their endangered status, and in some cases extinction (Sillero-Zubiri & Laurenson 2001, Ceballos et al. 2005, Schipper et al. 2008, Karanth & Chellam, 2009, Karanth et al. 2010). Recent assessments of the conservation status indicate alarming rates of population decline for many carnivores at a global scale (Ceballos et al. 2005, Schipper et al. 2008, Karanth & Chellam, 2009, Wolf and Ripple 2017). Specifically, the families *Felidae, Canidae* and *Ursidae* are under severe threat across the globe (Ceballos et al. 2005, Schipper et al. 2008, Karanth et al. 2010, Wolf and Ripple 2017).

Leopard (*Panthera pardus*) represents the most widely distributed and adaptable member of the family *Felidae*. The historical range of leopards spanned across nearly 35,000,000 km^2^ area covering all of sub-Saharan and North Africa, the Middle East and Asia Minor, South and Southeast Asia, and the Russian Far East (Uphyrkina et al. 2001, Jacobson et al. 2016). However, their current distribution and numbers have significantly decreased across the range due to habitat loss, prey depletion, conflict and poaching over the last century (Shepherd 2001, Banks & Newman 2004, Jacobson et al. 2016). Recent meta-analyses of leopard status and distribution suggest 48-67% range loss for the species in Africa and 83-87% in Asia (Jacobson et al. 2016), making them among the top ten large carnivore species most-affected by range contraction (Wolf and Ripple 2017). Out of the nine recognized leopard subspecies, three (*P. p. orientalis, P. p. nimr* and *P. p. melas*) are classified as Critically Endangered, while two (*P. p. saxicolor* and *P. p. kotiya*) are considered Endangered, and remaining four (*P. p. japonensis, P. p. delacouri, P. p. fusca* and *P. p. pardus*) are categorized as Near Threatened by IUCN (Stein et al. 2016). Despite continuously decreasing numbers and range, their ubiquitous presence across human habitations leads to misconceptions regarding their current abundance.

Among all the subspecies, the Indian leopard (*P. p. fusca*) retains the largest population size and range outside Africa (Jacobson et al. 2016). In the Indian subcontinent poaching and conflict are major threats to leopard populations (Edgaonkar & Chellam 1998, Athreya et al. 2010, Karanth et al. 2010, 2012, 2013a, b, Raza et al. 2012, Mondol et al. 2014). Leopards also frequently occur outside protected areas, increasing their vulnerability to conflict with humans (Karanth et al. 2009, Athreya et al. 2010, Karanth et al. 2010, 2012a,b, 2013, Naha et al. 2018). Unfortunately, there is still a paucity of information on their population and demography at regional and global scales. Much of our knowledge on leopard ecology and demography in the Indian subcontinent come from location-specific studies (Karanth & Sunquist 2000, Chauhan et al. 2005, Harihar et al. 2009, Wang and Macdonald 2009, Kalle et al. 2011, Grant 2012, Dutta et al. 2012a,b, Mondal et al. 2012, Borah et al. 2014, Selvan et al. 2014, Thapa et al. 2014, Pawar et al. 2019). In India, the latest estimate of leopards in the forested habitats of 14 tiger-inhabiting states is 7910 (SE 6566-9181) (Jhala et al. 2014). As leopards do survive in highly human populated and modified areas (Athreya et al 2013, 2014) this estimate is likely to be minimal and incomplete. Further, recent studies in the Indian subcontinent provide contradictory patterns of local population trends. For e.g., historical records and occupancy estimation models based on ecological data and field observations Karanth et al. (2010) estimated high local extinction probabilities of leopards across the subcontinent, and Athreya et al. (2010) reported higher rates of recent conflict incidences and related mortality at local scales. Other ecological (Harihar et al. 2011) as well as population genetic studies of demographic history (Dutta et al. 2012a) suggest stable or increased leopard populations at local scales. However, lack of detailed, systematic field data makes it difficult to generate accurate population estimates as well as demographic patterns at landscape scales.

In this paper, we used non-invasively collected faecal genetic data to assess leopard genetic variation, population structure and demographic history in the Indian subcontinent. More specifically, we investigated (1) extent of genetic variation in leopard that persists across the Indian subcontinent; (2) population structure of leopards at country scale; (3) the demographic history of leopards by assessing recent changes in population size and finally (4) compared the finding of genetic decline analyses with country wide ecological extinction probabilities. We interpreted our results in the context of local extinction probabilities as estimated in Karanth et al. (2010). We addressed these questions using genetic data generated using 13 polymorphic microsatellite loci from leopard faecal samples collected across different landscapes of India.

## Methods

### Research permissions and ethical considerations

All required permissions for our field surveys and biological sampling were provided by the Forest Departments of Uttarakhand (Permit no: 90/5-6), Uttar Pradesh (Permit no: 1127/23-2-12(G) and 1891/23-2-12) and Bihar (Permit no: Wildlife-589). Due to non-invasive nature of sampling, no ethical clearance was required for this study.

### Sampling

To detect population structure and past population demography it is important to obtain genetic samples from different leopard habitats all across the study area. In this study, we used leopard genetic data generated from non-invasive samples collected across the Indian subcontinent. We conducted extensive field surveys across the Indian part of Terai-Arc landscape (TAL) covering the north-Indian states of Uttarakhand, Uttar Pradesh and Bihar between 2016-2018. During the surveys, we opportunistically collected a total of 778 fresh large carnivore faecal samples. These samples were collected from both inside (n=469) and outside (n=309) protected areas from different parts of this landscape. In the field, the samples were judged as large carnivores based on several physical characteristics such as scrape marks, tracks, faecal diameter etc. All faecal samples were collected in wax paper and stored individually in sterile zip-lock bags and stored inside dry, dark boxes in the field for a maximum of two weeks period (Biswas et al. 2019). All samples were collected with GPS locations and were transferred to the laboratory and stored in −20°C freezers until further processing.

In addition to the north Indian samples collected in this study, we have also used leopard genetic data previously described in Mondol et al. (2014), representing mostly the Western Ghats and central Indian landscape. The data was earlier used in forensic analyses to assign seized leopard samples to their potential geographic origins in India (Mondol et al. 2014). Out of the 173 individual leopards described in the earlier study, we removed data from related individuals and samples with insufficient data (n=30) and used remaining 143 samples for analyses in this study. These samples were collected from the states of Kerala (n=5), Tamil Nadu (n=4), Karnataka (n=53), Andhra Pradesh (n=3), Madhya Pradesh (n=12), Maharashtra (n=46), Gujarat (n=2), Rajasthan (n=5), Himachal Pradesh (n=8), Jharkhand (n=1), West Bengal (n=2) and Assam (n=2), respectively. The sample locations across the Indian subcontinent those used in the final analyses are provided in Figure 1.

**Figure 1:**
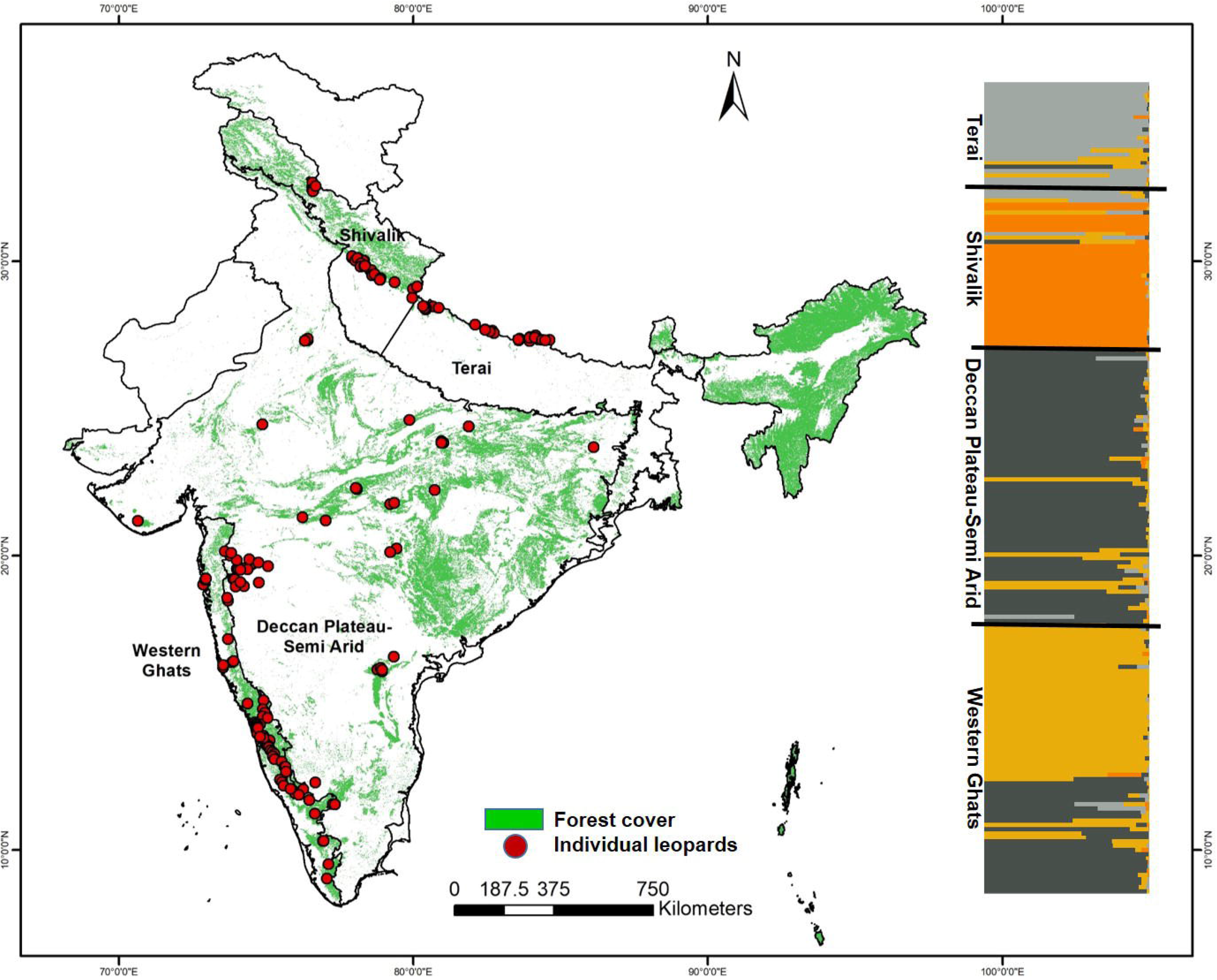
Forest cover map with leopard genetic sampling and population structure across the Indian subcontinent. The map shows the inferred biogeographic leopard habitats based on genetic cluster results found in this study. The STRUCTURE plot shows the partitioning of microsatellite genetic variation (K=4) based on 13 loci data. The cluster names correspond to the habitats across India subcontinent.

### DNA extraction, species and individual identification

For all field-collected faecal samples, DNA extraction was performed using protocols described in Biswas et al. (2019). In brief, each frozen faeces was thawed to room temperature and the upper layer was swabbed twice with Phosphate buffer saline (PBS) saturated sterile cotton applicators (HiMedia). The swabs were lysed with 30 μl of Proteinase K (20mg/ml) and 300 μl of ATL buffer (Qiagen Inc., Germany) overnight at 56°C, followed by Qiagen DNeasy tissue DNA kit extraction protocol. DNA was eluted twice in 100 μl preheated 1X TE buffer. For every set of samples, extraction negatives were included to monitor possible contaminations.

Species identification was performed using leopard-specific multiplex PCR assay described in Mondol et al. (2014) and Maroju et al. (2016). PCR reactions were done in 10 μl volumes containing 3.5 μl multiplex buffer mix (Qiagen Inc., Germany), 4 μM BSA, 0.2 μM primer mix and 3 μl of scat DNA with conditions including initial denaturation (95°C for 15 min); 40 cycles of denaturation (94°C for 30 s), annealing (T_a_ for 30 s) and extension (72°C for 35 s); followed by a final extension (72°C for 10 min). Negative controls were included to monitor possible contamination. Leopard faeces were identified by viewing species-specific bands of 130 and 190 bp (Mondol et al. 2014) and 277 bp (Maroju et al. 2016) in 2% agarose gel.

For individual identification, we used the same panel of 13 microsatellite loci previously used in Mondol et al. (2014) (Table 1). To generate comparable data with the samples used from earlier study by Mondol et al. (2014) we employed stringent laboratory protocols. All PCR amplifications were performed in 10 μl volumes containing 5 μl Qiagen multiplex PCR buffer mix (QIAGEN Inc., Germany), 0.2 μM labelled forward primer (Applied Biosystems, USA), 0.2 μM unlabelled reverse primer, 4 μM BSA and 3 μl of the faecal DNA extract. The reactions were performed in an ABI thermocycler with conditions including initial denaturation (94°C for 15 min); 45 cycles of denaturation (94°C for 30 sec), annealing (T_a_ for 30 sec) and extension (72°C for 30 sec); followed by final extension (72°C for 30 min). Multiple primers were multiplexed to reduce cost and save DNA (Table 1). PCR negatives were incorporated in all reaction setups to monitor possible contamination. The PCR products were analyzed using an automated ABI 3500XL Bioanalyzer with LIZ 500 size standard (Applied Biosystems, USA) and then alleles were scored with GENEMAPPER version 4.0 (Softgenetics Inc., USA). During data generation from field-collected samples we used one reference sample (genotyped for all loci) from the earlier study for genotyping. As the entire new data is generated along with the reference sample and the alleles were scored along with the reference genotypes, the new data (allele scores) was comparable with the earlier data for analyses.

**Table 1:** Genetic diversity and genotyping error details for the leopard samples collected across Terai Arc landscape (n=56) in this study. A total of 13 microsatellite loci were used. Data from these samples have been added to earlier leopard forensic data described in Mondol et al. (2014).

| Locus | Repeat length | N<sub>A</sub> | Allelic size range | H<sub>E</sub> | H<sub>O</sub> | Null Allele | Allelic dropout | False allele | HWE | Reference |
| --- | --- | --- | --- | --- | --- | --- | --- | --- | --- | --- |
| **FCA230** | 2 | 16 | 44 | 0.87 | 0.69 | 0.18 | 0.001 | 0.005 | Yes | Menotti-Raymond et al. (1999) |
| **FCA309** | 2 | 17 | 42 | 0.85 | 0.70 | 0.22 | 0.004 | 0.004 | Yes | Menotti-Raymond et al. (1999) |
| **FCA232** | 2 | 15 | 36 | 0.83 | 0.68 | 0.19 | 0.007 | 0.013 | Yes | Menotti-Raymond et al. (1999) |
| **FCA090** | 2 | 16 | 34 | 0.87 | 0.66 | 0.30 | 0.007 | 0.002 | Yes | Menotti-Raymond et al. (1999) |
| **FCA052** | 2 | 14 | 32 | 0.85 | 0.77 | 0.19 | 0.004 | 0.006 | Yes | Menotti-Raymond et al. (1999) |
| **FCA672** | 2 | 20 | 40 | 0.87 | 0.74 | 0.05 | 0.0 | 0.001 | Yes | Menotti-Raymond et al. (1999) |
| **FCA279** | 2 | 16 | 30 | 0.88 | 0.76 | 0.08 | 0.001 | 0.003 | Yes | Menotti-Raymond et al. (1999) |
| **FCA126** | 2 | 16 | 32 | 0.89 | 0.70 | 0.36 | 0.004 | 0.001 | Yes | Menotti-Raymond et al. (1999) |
| **msFCA391** | 4 | 10 | 36 | 0.86 | 0.64 | 0.19 | 0.009 | 0.007 | Yes | Mondol et al. (2011) |
| **msHDZ170** | 2 | 13 | 42 | 0.83 | 0.53 | 0.30 | 0.002 | 0.002 | Yes | Mondol et al. (2011) |
| **msFCA441** | 4 | 12 | 52 | 0.82 | 0.52 | 0.23 | 0.006 | 0.003 | Yes | Mondol et al. (2011) |
| **msFCA506** | 2 | 19 | 56 | 0.89 | 0.69 | 0.25 | 0.008 | 0 | Yes | Mondol et al. (2011) |
| **msFCA453** | 4 | 7 | 32 | 0.68 | 0.61 | 0.25 | 0.006 | 0.007 | Yes | Mondol et al. (2011) |
| **Mean (SD)** |  | **14.69**<br>**(3.41)** | **39.08**<br>**(7.71)** | **0.84**<br>**(0.05)** | **0.67**<br>**(0.07)** | **0.21** | **0.005** | **0.004** |  |  |

N<sub>A</sub> - No. of alleles, H<sub>E</sub> – Expected heterozygosity, H<sub>O</sub> – Observed heterozygosity, HWE – Hardy-Weinberg Equilibrium

To ensure good quality multi-locus genotypes from faecal samples, we followed a modified multiple-tube approach in combination with quality index analyses (Miquel et al. 2006) as described previously for leopards by Mondol et al. (2009, 2014). All faecal samples were amplified and genotyped four independent times for all the loci. Samples producing identical genotypes for at least three independent amplifications (or a quality index of 0.75 or more) for each loci were considered reliable and used for all further analysis, while the rest were discarded.

### Analysis

For each locus, we calculated average amplification success as the percent positive PCR (Broquet & Petit 2004) after four repeats across all samples. We quantified allelic dropout and false allele rates manually as the number of dropouts or false alleles over the total number of amplifications, respectively (Broquet & Petit, 2004), as well as using MICROCHECKER v 2.2.3. (Oosterhout et al. 2004). The false allele frequency is calculated for both homozygous and heterozygous genotypes as the ratio of the number of amplifications having one or more false alleles at a particular locus and the total number of amplifications while allele dropout rate (ADO) is calculated as the ratio between the observed number of amplifications having loss of one allele and the number of positive amplifications of the heterozygous individuals.

Post data quality assessment and finalization of consensus genotypes for all samples we selected only those samples with good quality data for at least nine or more loci (out of 13) for further analyses. We used the identity analysis module implemented in program CERVUS (Kalinowski, Taper & Marshall, 2007) to identify identical genotypes (or recaptures) by comparing data from all samples. All genetic recaptures were removed from the data set. GIMLET (Valiere, 2002) was used to calculate the PID_(sibs)_ for all the unique individuals. Following this, any allele having less than 10% frequency across all amplified samples were rechecked for allele confirmation. ARLEQUIN (Excoffier, Laval & Schneider, 2005) was used to determine Hardy Weinberg equilibrium and linkage disequilibrium for all the loci. Finally, to avoid the effects of related individuals in all analyses, we used program GENECLASS 2.0 (Piry et al. 2004) to select out related individuals in our samples.

To determine genetic structure of leopards across the Indian subcontinent we used a Bayesian clustering approach implemented in program STRUCTURE (Pritchard et al. 2000, Falush et al. 2003). We performed 10 independent analyses for each K values between one and ten, using 450,000 iterations and a burn-in of 50,000 assuming correlated allele frequencies. The optimal value of K was determined using STRUCTURE HARVESTER web version (Earl & vonHoldt, 2012). Subsequent summary statistics were calculated in ARLEQUIN 3.1 (Excoffier et al. 2005) and indices of overall genetic differentiation (pairwise F_st_) were estimated using GenAlEx version 6.5 (Peakall and Smouse 2012), dividing the leopard populations according to the STRUCTURE results across the Indian subcontinent. The divisions were based on Q-values (estimated proportions of ancestry) calculated in STRUCTURE, where we used Q> 0.75 as threshold for assigning individuals to a particular population (Mora et al. 2010). Additionally compression of expected heterozygosity (or G_st_) (Nei, 1987) between four leopard sub-populations was calculated in GenAlEx version 6.5 (Peakall and Smouse 2012).

### Demography analyses

Demographic analyses were performed with different genetic subpopulations of leopards based on the results from STRUCTURE analyses. We used a number of different approaches to detect past population demography for leopards. The first two qualitative approaches use summary statistics to detect population size changes, whereas the quantitative approach is a likelihood-based Bayesian algorithm. The summary statistic-based methods used were the Ewens, Watterson, Cornuet and Luikart method implemented in program BOTTLENECK (Cournet & Luikart 1996), and the Garza-Williamson index or M ratio (Garza & Williamson 2001) implemented in program ARLEQUIN 3.1. The quantitative Bayesian approach used was implemented in the program MSVAR 1.3 (Storz & Beaumont 2002).

#### a) The Ewens, Watterson, Cornuet and Luikart (EWCL) approach

This approach allows the detection of population size changes using two summary statistics of the allele frequency spectrum, number of alleles (N_A_) and expected heterozygosity (H_e_) across different mutational models. Simulations are performed to obtain the expected distribution of H_e_ for a demographically stable population under three mutation models: infinite allele model (IAM), single stepwise model (SMM) and two-phase model (TPM) and the values are then compared to the real data values. This method can detect departures from mutation-drift equilibrium and neutrality, which can be explained by any departure from the null model, including selection, population growth or decline. More importantly, consistent results from independent loci could be attributed to demographic events over selection. For simulations with TPM model, we used two different (5% and 30%) multi-step mutation events for leopards.

#### b) The Garza-Williamson index/M ratio approach

This approach allows the detection of population decline using two summary statistics of the allele frequency spectrum, number of alleles (N_A_) and the allelic size range. The basic principle behind this approach is in a reducing population, the expectation of the reduction of number of alleles is much higher than the reduction of allelic size range. Thus, the ratio between the number of alleles and the allelic size range is expected to be smaller in recently reduced populations than in equilibrium populations.

#### c) The Storz and Beaumont approach

This approach is an extension of Beaumont’s approach (Beaumont 1999) that assumes a stable population of size N_1_ started to change (either decrease or increase) T_a_ generations ago to the current population size N_0_. This change in the population size is assumed to be at an exponential scale under stepwise mutation model (SMM), at a rate y=2N_0_m, where m is the mutation rate per locus per generation. This Bayesian approach uses the information from the full allelic distribution in a coalescent framework to estimate the posterior probability distribution, allowing quantification of effective population sizes N_0_ and N_1_, rather than their ratio (as in Beaumont 1999) along with T, time since the population change. In this approach, prior distributions for N_0_, N_1_, T and μ (mutation rate) are assumed to be log normal. The mean and the standard deviations of these prior log normal distributions are drawn from prior (or hyperpriors) distributions. A Markov Chain Monte Carlo (MCMC) algorithm is used to generate samples from the posterior distribution of these parameters. We used wide uninformative priors to perform multiple runs for this approach (Supplementary Table 1). For minimal effect towards the posterior distributions variances for the prior distributions were kept large. A total number of 2 million iterations were performed for each run.

The generation time for leopards are known to be about 4-5 years (Dutta et al. 2012b) and we used a five-year generation time for all analyses.

### Estimation of leopard extinction probability

To understand extinction probability across various biogeographic zones of India we analysed patterns and determinants of leopard occurrence as described in Karanth et al. (2009, 2010). We applied a grid-based approach to determine current distribution patterns for leopards, where the selection of grids was based on prior information of leopard presence. This involved collating presence-absence information from more than 100 Indian wildlife experts along with historical information of leopard presence involving hunting locations and other taxidermy and museum records. Each grid cell was an average of 2818 km^2^ in size and we used data from 1229 grid cells covering 3,46,3322 km^2^ area of the Indian subcontinent. This study applied occupancy modelling to examine the influence of ecological and social covariates on patterns of leopard occupancy. We used a maximum likelihood approach for leopard occupancy in PRESENCE. V.2.0 program (Hines 2006). Covariates likely to influence leopard distribution modelled included presence and extent of protected areas, land cover-land use characteristics, human cultural tolerance and population density. Data for protected areas was retrieved from the World Database on protected areas (www.unep-wcmc.org) and topographic maps. Land cover-land use data were derived from Global Land Cover Facility (2000) and further refined based on Roy et al. (2006) and Joshi et al. (2006). A human tolerance index that characterized different Indian states from most to least tolerant was developed based on knowledge about society-culture, law enforcement, hunting patterns and prior field experiences (for details see Karanth et al. 2009, 2010). Human population density data were derived from LandScan Global Population Data 2000 (www.ornl.gov/gist). Based on existing information on species’ ecology we predicted higher occupancy in protected areas, deciduous-grass-scrub land cover types and lower occupancy in less tolerant states and highly populated areas because of direct completion for food and space (Rangarajan 2001). We performed pair-wise correlation tests to screen variables for multicollinearity. The occupancy approach accounts for non-detection of species during surveys and inability to survey some sites (see Karanth et al. 2009, 2010 for additional details). The probability of extinction was calculated as (1-probability of occurrence). We derived leopard extinction probabilities for three separate major landscapes (Western Ghats, Central India and North India) as these regions strongly represented our genetic sampling. These extinction probabilities were compared to the genetically derived estimates.

## Results

### Individual identification of leopards from north Indian landscape

Of the 778 large carnivore faecal samples collected from TAL, we identified 195 faeces to be of leopard origin (25%) using species-specific PCR assays (Mondol et al. 2014; Maroju et al. 2016). In addition, 457 samples were ascertained to be of tiger (59%) and remaining 126 faecal samples did not produce any result (16%). We amplified 13 microsatellite loci panel on these 195 genetically confirmed leopard faecal samples, and after data validation through multiple repeats generated seven or more loci data from 65 faecal DNA. Subsequently, we identified 56 unique leopard individuals from the 65 samples, whereas nine individuals were ascertained as ‘genetic recaptures’. The mean allelic dropout rate for these loci was found to be 0.05, whereas mean false allele rate for all the 13 loci was 0.04., indicating this 13 loci panel has low genotyping error rates. Amplification success ranged between 41% to 100% from leopard faecal DNA. None of the loci were found to deviate from the Hardy-Weinberg equilibrium and there were no evidences for strong linkage disequilibrium between any pair of loci. Cumulative PID_sibs_ and PID_unbiased_ values were found to be 3.91*10^−6^ and 2.73*10^−16^, respectively, indicating a strong statistical support for unambiguous individual identification. Summary statistics for these samples collected across Terai-Arc landscape is provided in Table 1. We identified 26, 21 and nine unique leopard individuals from the states of Uttarakhand, Uttar Pradesh and Bihar, respectively. As the data generated from north India is comparable to the earlier data, we added this 56 unique leopard data to 143 individual genotypes described in Mondol et al. (2014), and overall 199 unique unrelated leopards were used in subsequent population structure, genetic variation and demography analyses.

### Leopard population structure and genetic variation across India

Our sampling strategy targeted country-wise leopard populations to assess population structure and genetic variation. From 199 final unique leopard genotypes we removed four samples representing the eastern and northeast India (n=2 from the states of West Bengal and Assam each, respectively) from further analyses as they represented inadequate sampling from these regions. Genetic clustering analysis using 13 microsatellite data from the remaining 195 wild leopard individuals showed four distinct genetic subpopulations (K=4, see Supplementary Figure 1), as presented in Figure 1. Majority of the samples showed respective group-specific ancestry, with Western Ghats samples representing the first group (henceforth WG, n=65), the Deccan Plateau-Semi Arid region forming the second cluster (henceforth DP-SA, n=66), the samples from Shivalik region covering parts of Himalaya and western parts of upper Gangetic plains making the third group (henceforth SR, n=38), and finally samples from the Terai region covering eastern part of upper and western part of the lower Gangetic Plains samples forming the fourth cluster (henceforth TR, n=26), respectively (Figure 1). However, small number of samples (n=18) distributed among the four subpopulations showed mixed ancestry. Subsequent analyses revealed that these leopard subpopulations are genetically differentiated (F_st_ and G_st_) at low, but significant levels (Table 2) for all four populations. The F_st_ value among these populations ranged between 0.028-0.115, whereas the G_st_ value between 0.023-0.104 (Table 2).

**Table 2:** Genetic differentiation (pairwise F_st_ and G_st_) for four leopard subpopulations in the Indian subcontinent. The upper diagnonal presents the pairwise Gst values whereas the lower diagnonal presents the pairwise Fst values.

|  | Western Ghats | Deccan Plateau-Semi Arid | Shivalik | Terai |
| --- | --- | --- | --- | --- |
| Western Ghats | -- | 0.023* | 0.039* | 0.091* |
| Deccan-Semi Arid | 0.028* | -- | 0.045* | 0.104* |
| Shivalik | 0.048* | 0.05* | -- | 0.073* |
| Terai | 0.103* | 0.115* | 0.089* | -- |

\* p value =0.001

Analyses and with 13 microsatellite loci among the four genetic subpopulations showed a higher mean number of alleles (NA_WG_= 11.77 (S.D. 3.85), NA_DP-SA_=10.46 (S.D. 2.71)) and observed heterozygosity (H_oWG_=0.81 (S.D. 0.08), H_oDP-SA_=0.8 (S.D. 0.08)) in Western Ghats and Deccan Plateau-Semi Arid subpopulations, when compared with samples from Shivalik and Terai region subpopulations (NA_SR_=08.46 (S.D. 2.41), NA_TR_=05.00 (S.D. 1.84) and H_oSR_=0.40 (S.D. 0.14), H_oTR_=0.36 (S.D. 0.28), respectively) (see Table 3 for details). However, the allelic size range values were similar in all populations (Table 3).

**Table 3:** Subpopulation-wise summary statistics (based on 13 microsatellite loci) for Indian leopards.

|  | Western Ghats (n=65) |  |  |  | Deccan Plateau-Semi Arid (n=66) |  |  |  | Shivalik (n=38) |  |  |  | Terai (n=26) |  |  |  |
| --- | --- | --- | --- | --- | --- | --- | --- | --- | --- | --- | --- | --- | --- | --- | --- | --- |
| Locus | N<sub>A</sub> | ASR | H<sub>E</sub> | H<sub>O</sub> | N<sub>A</sub> | ASR | H<sub>E</sub> | H<sub>O</sub> | N<sub>A</sub> | ASR | H<sub>E</sub> | H<sub>O</sub> | N<sub>A</sub> | ASR | H<sub>E</sub> | H<sub>O</sub> |
| **FCA230** | 13 | 36 | 0.88 | 0.86 | 10 | 22 | 0.78 | 0.80 | 08 | 24 | 0.83 | 0.23 | 05 | 26 | 0.52 | 0.65 |
| **FCA309** | 17 | 42 | 0.90 | 0.87 | 11 | 30 | 0.78 | 0.81 | 08 | 18 | 0.81 | 0.46 | 06 | 10 | 0.82 | 0.32 |
| **FCA232** | 13 | 36 | 0.85 | 0.84 | 09 | 18 | 0.68 | 0.72 | 09 | 26 | 0.78 | 0.42 | 07 | 26 | 0.78 | 0.46 |
| **FCA090** | 14 | 30 | 0.85 | 0.84 | 08 | 18 | 0.78 | 0.87 | 09 | 30 | 0.86 | 0.36 | 02 | 10 | 0.47 | 0.00 |
| **FCA052** | 12 | 32 | 0.84 | 0.89 | 11 | 22 | 0.82 | 0.84 | 08 | 20 | 0.83 | 0.48 | 06 | 22 | 0.84 | 0.43 |
| **FCA672** | 19 | 40 | 0.90 | 0.89 | 10 | 26 | 0.82 | 0.75 | 06 | 16 | 0.65 | 0.50 | 07 | 20 | 0.64 | 0.77 |
| **FCA279** | 11 | 26 | 0.81 | 0.75 | 14 | 26 | 0.87 | 0.81 | 15 | 28 | 0.90 | 0.67 | 06 | 18 | 0.78 | 0.92 |
| **FCA126** | 14 | 26 | 0.87 | 0.83 | 13 | 30 | 0.88 | 0.89 | 09 | 22 | 0.76 | 0.18 | 04 | 12 | 0.11 | 0.74 |
| **msFCA391** | 07 | 28 | 0.83 | 0.67 | 08 | 32 | 0.81 | 0.81 | 07 | 24 | 0.78 | 0.57 | 07 | 32 | 0.75 | 0.13 |
| **msHDZ170** | 09 | 20 | 0.84 | 0.79 | 10 | 22 | 0.75 | 0.88 | 10 | 36 | 0.76 | 0.16 | 02 | 02 | 0.29 | 0.0 |
| **msFCA441** | 07 | 36 | 0.75 | 0.79 | 08 | 28 | 0.65 | 0.55 | 08 | 40 | 0.82 | 0.38 | 05 | 28 | 0.66 | 0.40 |
| **msFCA506** | 12 | 32 | 0.86 | 0.90 | 17 | 56 | 0.83 | 0.82 | 09 | 24 | 0.83 | 0.33 | 06 | 22 | 0.79 | 0.40 |
| **msFCA453** | 05 | 20 | 0.63 | 0.67 | 07 | 32 | 0.69 | 0.80 | 04 | 20 | 0.65 | 0.36 | 02 | 16 | 0.43 | 0.14 |
| **Mean (SD)** | **11.77**<br>**(3.85)** | **31.08**<br>**(6.69)** | **0.83**<br>**(0.07)** | **0.81**<br>**(0.08)** | **10.46**<br>**(2.71)** | **27.85**<br>**(9.36)** | **0.78**<br>**(0.07)** | **0.80**<br>**(0.08)** | **08.46**<br>**(2.41)** | **25.23**<br>**(6.64)** | **0.79**<br>**(0.07)** | **0.40**<br>**(0.14)** | **05.00**<br>**(1.84)** | **18.77**<br>**(8.21)** | **0.65**<br>**(0.17)** | **0.36**<br>**(0.28)** |

N<sub>A</sub> - No. of alleles, ASR- Allelic size range, H<sub>E</sub> – Expected heterozygosity, H<sub>O</sub> – Observed heterozygosity,

### Detection of demographic change

We used the microsatellite data to investigate signals of demographic changes in each of the four leopard genetic subpopulations across the subcontinent. Both of the qualitative approaches, the EWCL and the M-ratio methods indicate signatures of population bottleneck. The EWCL approach implemented in the program BOTTLENECK shows 8-10 loci with heterozygote excess depending on the mutation models used, suggesting a loss of rare alleles through population decline for all four subpopulations. Similarly the M-ratio approach also shows a low ratio between number of alleles (N_A_) and the allelic size range in all four populations (M-ratio_WG_-0.37 (S.D. 0.09); M-ratio_DP-SA_-0.38 (S.D. 0.09); M-ratio_SR_-0.33 (S.D. 0.09); M-ratio_TR_-0.29 (S.D. 0.15)), indicating signatures of population bottleneck. However, both of these approaches cannot quantify the extent and timing of the bottleneck events. We used the Storz and Beaumont approach for quantification and dating of any such events through coalescent simulations. Models with exponential decline scenarios show consistently that the posterior distributions for log (N0) is always lower than log (N1) for all four subpopulations, indicating population decline for leopards across the subcontinent (Table 4 and Figure 2). Further quantification revealed that the current effective size is varyingly low (12-25%) than the historical effective size, with Western Ghats, Deccan Plateau-Semi Arid, Shivalik and Terai regions losing approximately 75%, 90%, 90% and 88% of their leopard population, respectively (Table 4 and Figure 2).

**Figure 2:**
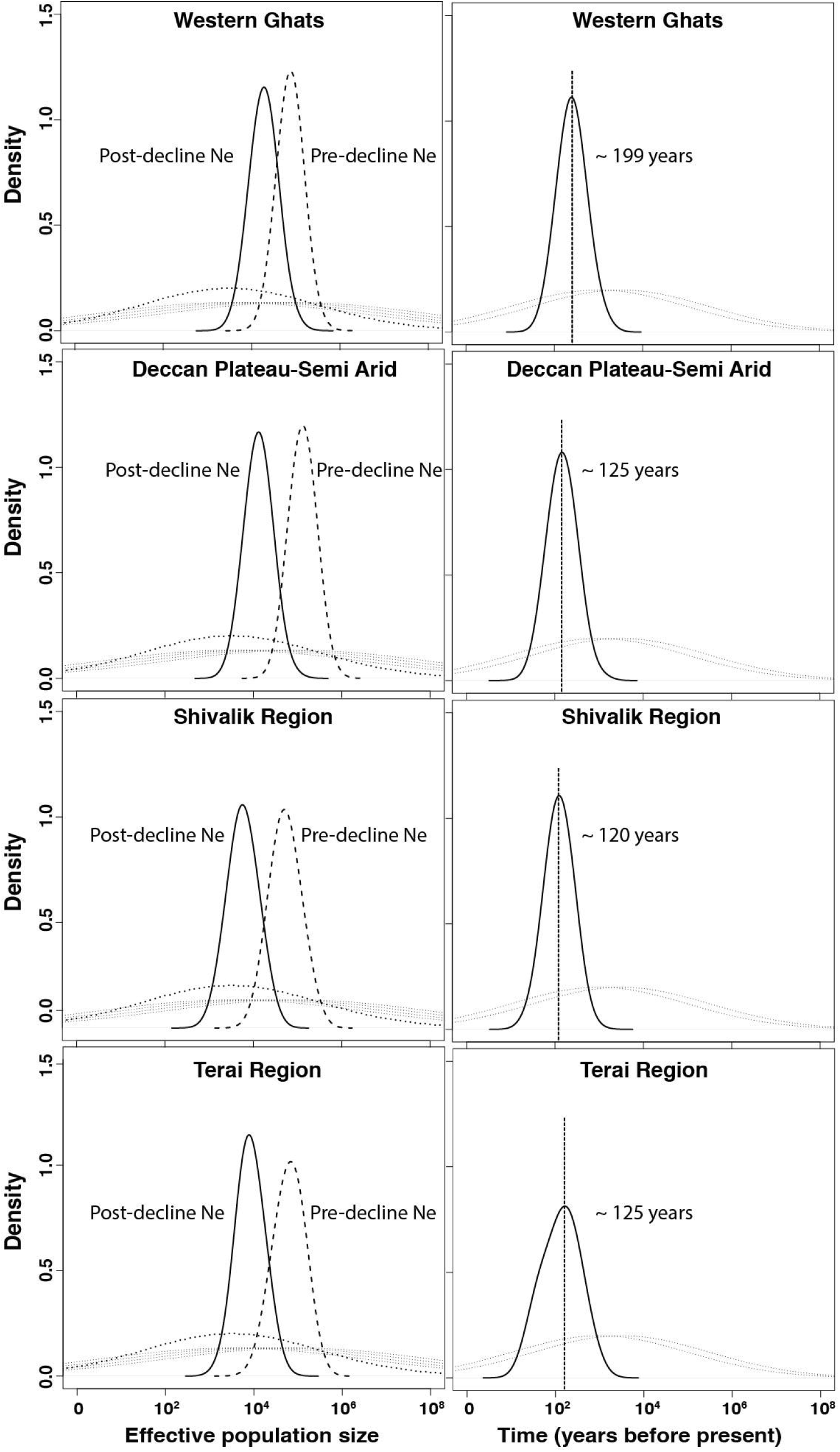
Demographic history of different genetic subpopulations of Indian leopards (*Panthera pardus fusca*). The left panel shows the posterior distributions for population size changes based on coalescent simulations for different leopard subpopulations using 13 microsatellite loci and the Storz and Beaumont approach. The dashed and solid lines represent posterior distributions for ancestral and present effective population sizes, respectively. The right panel presents the posterior distributions for the time since the population decline started for different leopard subpopulations. The distributions have median values (shown as vertical lines) ranging from ~120-199 years. In both cases priors are shown by fine dotted lines.

**Table 4:** Comparison of different demographic decline analyses results for different subpopulations of leopards across India.

| Method | Analysis type | Demographic signal |  |  |  |  |
| --- | --- | --- | --- | --- | --- | --- |
|  |  | Model | Western Ghats | Deccan Plateau-Semi Arid | Shivalik | Terai |
| Bottleneck | Qualitative | IAM | Heterozygosity excess for 13 loci | Heterozygosity excess for 10 loci | Heterozygosity excess for 12 loci | Heterozygosity excess for 11 loci |
|  |  | SMM | Heterozygosity excess for 01 loci | Heterozygosity excess for 02 loci | Heterozygosity excess for 06 loci | Heterozygosity excess for 08 loci |
|  |  | TPM | Heterozygosity excess for 07 loci | Heterozygosity excess for 07 loci | Heterozygosity excess for 09 loci | Heterozygosity excess for 10 loci |
| M ratio |  |  | 0.37 (SD 0.09) | 0.38 (SD 0.09) | 0.33 (SD 0.09) | 0.29 (SD 0.15) |
| Storz-Beaumont method | Quantitative |  | Decline- 75%<br>Time- ~200 years ago | Decline- 90%<br>Time- ~125 years ago | Decline- 90%<br>Time- ~125 years ago | Decline- 88%<br>Time- ~120 years ago |

This approach also allowed us to date the population collapse by providing a posterior distribution for the time at which the decline started. Our analyses revealed distributions that suggested recent time of declines in all four populations of leopards (Table 4, Figure 2). The north Indian subpopulations (Shivalik and Terai) and the Deccan Plateau-Semi Arid population showed the most recent decline occurred about 120-125 years before present, respectively. However, the Western Ghats population indicated potential decline around 200 years ago (Table 4 and Figure 2).

### Leopard occurrence and distribution

We examined the factors influencing leopard distribution at a countrywide scale, where the top ranked model incorporating 28 covariates suggested a wide distribution of habitat types (described in Karanth et al. 2009, 2010). The model also indicated a positive influence of protected areas and negative influence of higher human population densities and higher cultural tolerance of people (details in Karanth et al. 2009). Areas with cultivated land, barren areas, deciduous forests and rural-urban were strongly associated with higher leopard occurrence. Naive estimated occupancy was 0.52, whereas model estimated probability of occupancy was significantly higher at 0.68, suggesting that leopards are still widely distributed (Figure 3) in India compared to most other large mammals (as suggested in Karanth et al. (2010)). When compared among the overall three major sub-regions (North India (NI), Deccan Plateau-Semi Arid and Western Ghats), we find that average estimated occurrence was lowest in the North India (Psi_NI_=0.63, Range: 0.05-1.00, Standard error=0.01, Number of cells=384) compared to Western Ghats (Psi_WG_=0.83, Range: 0.23-1.00, Standard error=0.02, Number of cells=90) and Deccan Plateau-Semi Arid (Psi_DP-SA_=0.79, Range:0.25-1.00, Standard error=0.005, Number of cells=818). Overall, average estimated Psi was 0.74 (Standard error=0.006, Number of cells=1292).

**Figure 3:**
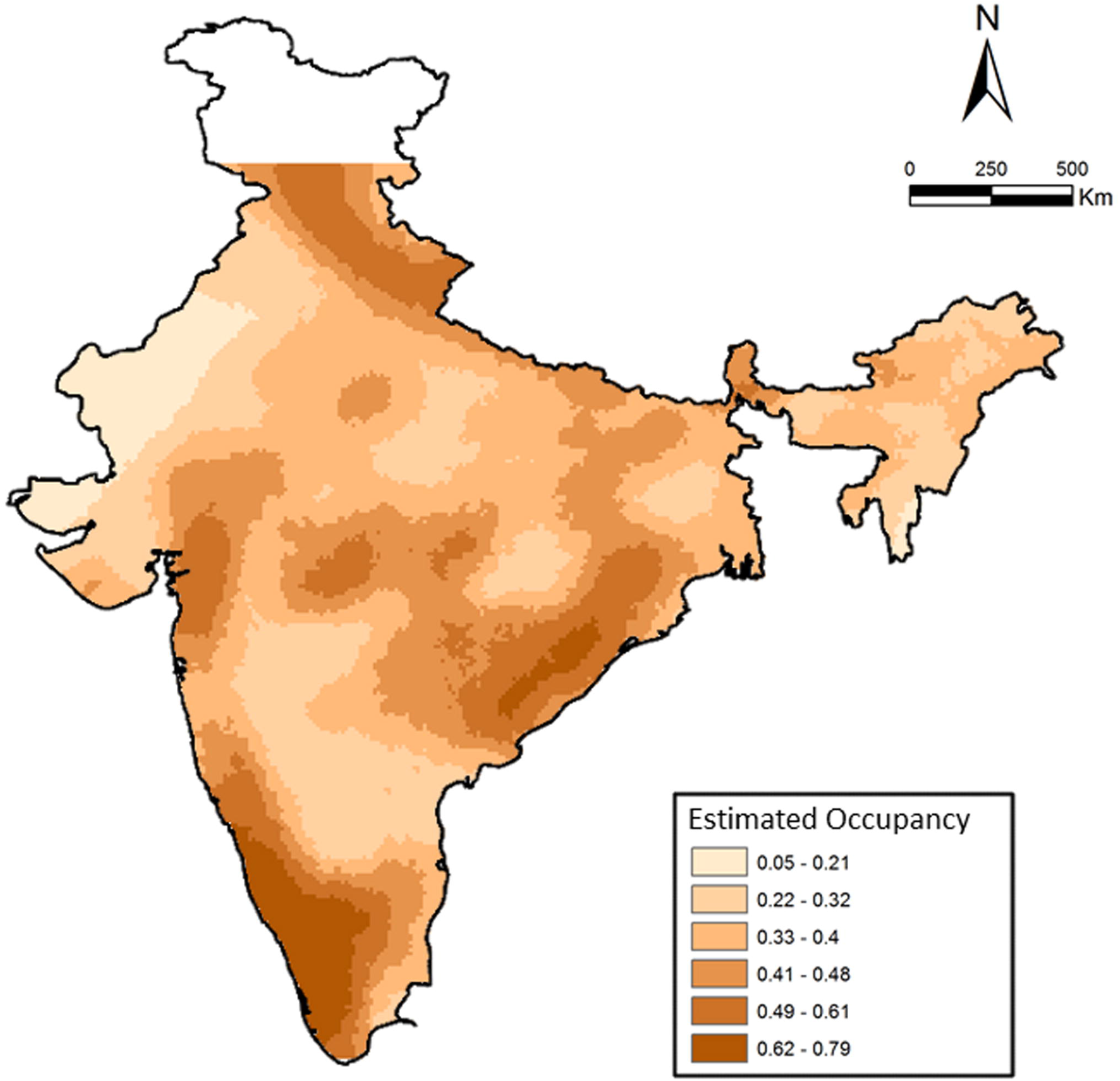
Patterns of leopard occurrence in India based on the analysis of questionnaire surveys. The map shows a gradient of estimated cell-wise occupancy probabilities created through spatial kriging.

## Discussion

To the best of our knowledge, this is probably the first and most exhaustive country level sampling based study on leopard population genetics and demographic patterns in the Indian subcontinent. Except the eastern and northeast Indian landscape, where our sampling intensity was less all other regions are well covered in this study. Our genetic analyses with microsatellite data collected across the subcontinent reveal four genetic subpopulations of leopards in India: the Western Ghats, Deccan Plateau-Semi Arid landscape, hill region of north India (Shivalik) and Terai or flat region of north India. While there was some amount of mixed genetic signal across different genetic subpopulations, they were clearly separated as different clusters (Figure 1). These genetic clusters mostly correspond to respective biogeographic zones of India, with Western Ghats and combination of Deccan Plateau-Semi Arid forms two clusters, whereas the north Indian clusters of Shivalik and Terai are parts of the Himalayan and Gangetic Plains zones, respectively. It is possible that these genetic clusters were formed due to the species distribution across various habitat types in different biogeographic zones across Indian subcontinent. For example, earlier study has reported difference in leopard occupancy in ‘Bhabar’ habitats of Shivalik (high abundance) and flat Terai region (low abundance) due to presence of socially dominant tigers and the absence of rugged escape terrain for leopards (Johnsingh et al. 2004). Such habitat heterogeneities might have resulted in developing genetic differences across these landscapes.

Overall, these four subpopulations were genetically differentiated by low, but significant F_st_ and G_st_ values across all comparisons (F_st_ and G_st_ values ranging from 0.028-0.115 and 0.023-0.104, respectively, see Table 2). Previous studies on tigers (Mondol et al. 2009, 2013, Kolipakam et al. 2019, subcontinent scale) as well as leopards (Dutta et al. 2012b, central Indian landscape) suggested long-distance movement as a potential cause for low genetic differentiation between populations. Leopards are more widely distributed than other sympatric large carnivores (tiger, lion) in the Indian subcontinent due to their general adaptability and wider diet spectrum (Seidensticker et al. 1990, Edgaonkar & Chellam 1998, Athreya et al. 2010), and disperse long distances (Sunquist 1983, Bailey 1993) often through human habitats. In addition, human-leopard conflict driven translocation is common in many parts in India (Athreya et al. 2010). Together, natural dispersal abilities and ‘human mediated gene flow’ because of translocation might be responsible for the low genetic differentiation among leopard subpopulations across the subcontinent. Earlier work in central Indian landscape (Dutta et al. 2012b) suggested a reduction in gene flow at recent times due to habitat destruction, but our study did not focus to answer such questions. Future studies should focus on using historical samples (e.g. museum skins, bones etc.) to assess any possible change in gene flow among leopard populations (For e.g. see Martinez-Cruz et al. 2007, Valdiosera et al. 2008, Lorenzen et al. 2011, Mondol et al. 2013) at subpopulation levels across the country.

However, our demography analyses with genetic data indicate strong decline in leopard population size across all four genetic subpopulations. Results with both qualitative (bottleneck and M-ratio approach) as well as quantitative (Storz and Beaumont approach) analyses revealed strong, but varying signals of population decline in all four subpopulations (Table 4). The Deccan Plateau-Semi Arid, Shivalik and Terai subpopulations show 90%, 90% and 88% decline in population size, respectively, whereas the Western Ghats population show relatively less (75%) decline in population size (Table 4). This pattern is possible as the Western Ghats still retains possibly the largest contiguous forested landscape with multiple interconnected protected area landscape, whereas the other regions have lot of human activities, possibly affecting leopard populations living in them. Further, the ecological data based occupancy analysis showed extinction probabilities of 0.37, 0.21 and 0.17 for North India, Deccan Plateau-Semi Arid and Western Ghats landscape, respectively (Table 4). This is not surprising as throughout their distribution leopards are closely associated with human population, making them vulnerable to conflict and poaching (Gavashelishvili & Lukarevskiy 2008, Athreya et al. 2010, Balme et al. 2010). While there is a discrepancy between the magnitudes of decline based on genetic and ecological models, it is possible that such pattern is because the ecological methods are more spatial, and inference is based on how much area leopards occupied in the past and how this has changed. However, if densities of leopards were high in the past, loss of even small habitats could result in the loss of many individuals. Since no quantitative comparisons for leopard density between the Western Ghats, Deccan Plateau-Semi Arid and North India is currently available, we cannot conclusively infer the former, but further research should investigate leopard densities and their temporal changes across the country. Finally, this decline pattern also roughly corroborates with 83-87% leopard range loss in Asia, indicating that habitat loss is possibly leading to population decline.

The magnitude of decline for leopards found in this study is contrasting to some of the earlier leopard studies in the subcontinent (for e.g. ecological work by Harihar et al. 2011, and genetic work by Dutta et al. 2012a) and Africa (Spong et al. 2000b), which suggest stable or increasing local leopard population trends. This is certainly possible as many of these studies were conducted inside protected areas, where leopard population dynamics depends on presence/absence of other large carnivores (tiger, dhole etc.) and other ecological factors. However, only 11% of Indian leopard distribution is within protected area network (Jacobson et al 2016), and our sampling at subcontinent scale is thus probably indicating the decline patterns at much larger scale. Nevertheless, this pattern of population decline is consistent with many other endangered species in the Asian region (for e.g. tiger-98% decline, Mondol et al. 2009; giant panda-90% decline, Zhu et al. 2010; orangutan-95% decline, Goossens et al. 2006; red panda-98% decline, Hu et al. 2011; Prezwalski’s gazelle-99% decline, Yang and Jiang 2011 etc.) as well as top carnivore species across the globe (for e.g. Finnish wolf-92% decline, Aspi et al. 2006; Otter-75% decline, Hajkova et al. 2007; Golden eagle-94% decline, Bourke et al. 2010; African wild dog-70% decline, Marsden et al. 2012; Fisher-90% decline, Tucker et al. 2012 etc.).

Another important finding is the relatively recent timing of decline for all the leopard subpopulations in the subcontinent. Our results suggest median leopard decline timing between 120-200 years across four genetic subpopulations (Table 4). Except Western Ghats (decline timing of ~200 years), all other subpopulations indicate much recent population decline (Central India-Deccan Plateau ~125 years, Shivalik ~120 years and Terai ~125 years). When compared with other sympatric, endangered species in the subcontinent (for e.g. tiger decline ~200 years ago; Mondol et al. 2009) or in the region (for e.g. Orangutan- ~210 years, Goossens et al. 2006; Giant panda- ~250 years, Zhu et al. 2010) this still seems to be much recent event. Other wide-ranging carnivores across the globe (for e.g. European wolves Aspi et al. 2006; African wild dog-Marsden et al. 2012; Eurasian badgers-Franz et al. 2014 etc.) too faced much longer decline period than leopards. One plausible explanation could be recent increase in leopard-human conflict (Athreya et al. 2010) and poaching intensity due to large demand of leopard body parts in the illegal wildlife markets (Raza et al. 2012; WPSI 2014). Historically, major leopard hunting events had been recorded across the Indian subcontinent during Mughal times (about 500-600 years ago), followed by colonial British bounty-hunting rule between 1850-1920 (Rangarajan 2006). However, large-scale landscape modification and fragmentation by human during the last century (central India-Rangarajan 1999, north India-Rangarajan 2006), coupled with poaching and conflict has possibly resulted in much recent loss of leopard populations across the country. Apart from sporadic information, we lack comprehensive data, both at historical as well as modern scales to investigate the true causes behind such patterns of differential population decline timing. For e.g., Dutta et al. (2012b) showed that during last three centuries severe changes in landscape characteristics (Settlement, villages, wild lands, human density) have occurred in the central Indian leopard habitats. However, we lack information on hunting and conflict levels from these regions. Future efforts should generate this important information to get an idea of the scenarios leading to such strong decline in a wide-ranging species like leopard. Finally, it is important to point out that in this study we have only explored relatively simple decline scenarios, and future studies should evaluate more detailed, computationally intensive demographic analyses with genome wide molecular markers (For e.g. see Franz et al. 2014, Nater et al. 2015) for better understanding of complex decline scenarios.

Another important aspect of the results from this study is that despite severe decline (Table 4) and small, but significant population structure (Figure 1B, Table 2) leopards still retain high genetic variation in the Indian subcontinent. We found that leopard genetic variation across four genetic subpopulations is similar and comparable to Africa (Spong et al. 2000b, Uphyrkina et al. 2001), and higher than Arabian (Ilani 1981, Perez et al. 2006) and Amur leopards (Uphyrkina et al. 2001, 2002, Sugimoto et al. 2014). The higher levels of variation could possibly be attributed to still relatively large population size, high pre-bottleneck genetic variation and potential historical gene flow across large landscapes.

### Conclusion

While leopards are relatively easier to study than other sympatric carnivores like tigers due to their ubiquitous presence, studies on their ecology are limited. In fact, due to their broad geographic distribution, leopard populations are perceived to be stable, with current IUCN Red List status of ‘vulnerable’. However, both historical records and recent conflict with humans suggest potentially declining population trends. Using genetic data, we reveal a strong signal of population decline (between 75-90%) across different habitats in the Indian subcontinent over the last 120-200 years. Our results are interesting because we demonstrate population decline in a wide-ranging and, commonly perceived as locally abundant species like the leopard, suggesting that leopards demand similar conservation attention like tigers in India. While we are unable to corroborate these population decline patterns with leopard census data, our results suggest that it will be important to generate such ecological abundance estimates for leopard populations in the near future. This work also emphasizes the importance of similar work on wide-ranging species, as it is possible that other species like the leopard may show population declines, especially in the context of the Anthropocene.

## Acknowledgement

We acknowledge the Director, Dean and Nodal Officer of Wildlife Forensics and Conservation Genetics Cell of Wildlife Institute of India for their support in this work. Our sincere thanks to Forest Departments of Uttarakhand, Uttar Pradesh and Bihar for research permits. We thank Dr. Uma Ramakrishnan of National Centre for Biological Sciences for providing reference leopard samples. Mr A. Madhanraj has provided critical support in genotyping facility and Mr. Debanjan Sarkar helped with GIS in the laboratory. We thank all the lab members of Wildlife Forensic and Conservation Genetics cell and especially Meercat lab for productive discussions and valuable comments. We also thank Dr. S. K. Gupta and Dr. S. P. Goyal for logistic support; our field assistants Annu, Bura, Abbhi, Ranjhu and Imam for their effort in the field. This research was funded by Wildlife Conservation Trust-Panthera Global Cat Alliance Grants and Department of Science and Technology, Government of India grant no EMR/2014/000982. Samrat Mondol was supported by the Department of Science and Technology INSPIRE Faculty Award (No.IFA12-LSBM-47).

## References

Aspi J, Roininen E, Ruokonen M, Kojola I, Vila C. 2006. Genetic diversity, population structure, effective population size and demographic history of the Finnish wolf population. Molecular Ecology 15:1561–1576. DOI: 10.1111/j.1365-294X.2006.02877.x.

Athreya V, Odden M, Linnell JDC, Karanth KU. 2010. Translocation as a Tool for Mitigating Conflict with Leopards in Human-Dominated Landscapes of India. Conservation Biology 25:133–141. DOI: 10.1111/j.1523-1739.2010.01599.x.

Balme GA, Slotow R, Hunter LTB. 2009. Impact of conservation interventions on the dynamics and persistence of a persecuted leopard (Panthera pardus) population. Biological Conservation 142:2681–2690. DOI: 10.1016/j.biocon.2009.06.020.

Banks D, Newman J. 2004. The Tiger Skin Trail.:21.

Beaumont MA. 1999. Detecting population expansion and decline using microsatellites. Genetics 153:2013–2029.

Biswas S, Bhatt S, Paul S, Modi S, Ghosh T, Habib B, Nigam P, Talukdar G, Pandav B, Mondol S. 2019. A practive faeces collection protocol for multidisciplinary research in wildlife science. Current Science 116:1878–1885. DOI: 10.18520/cs/v116/i11/1878-1885.

Borah J, Sharma T, Das D, Rabha N, Kakati N, Basumatary A, Ahmed MF, Vattakaven J. 2014. Abundance and density estimates for common leopard Panthera pardus and clouded leopard Neofelis nebulosa in Manas National Park, Assam, India. Oryx 48:149–155. DOI: 10.1017/S0030605312000373.

Bourke BP, Frantz AC, Lavers CP, Davison A, Dawson DA, Burke TA. 2010. Genetic signatures of population change in the British golden eagle (Aquila chrysaetos). Conservation Genetics 11:1837–1846. DOI: 10.1007/s10592-010-0076-x.

Broquet T, Petit E. 2004. Quantifying genotyping errors in noninvasive population genetics. Molecular Ecology 13:3601–3608. DOI: 10.1111/j.1365-294X.2004.02352.x.

Ceballos G, Ehrlich PR, Sobero J, Salazar I, Fay JP. 2005. Global Mammal Conservation: What Must We Manage? Science 309:603–607. DOI: 10.1126/science.1110063.

Dutta T, Sharma S, Maldonado JE, Wood TC, Panwar HS, Seidensticker J. 2013. Gene flow and demographic history of leopards (Panthera pardus) in the central Indian highlands. Evolutionary Applications 6:949–959. DOI: 10.1111/eva.12078.

Dutta T, Sharma S, Maldonado JE, Wood TC, Seidensticker J. 2012. A reliable method for individual identification and gender determination of wild leopards (Panthera pardus fusca) using non-invasive samples. Conservation Genetics Resources 4:665–667. DOI: 10.1007/s12686-012-9618-5.

Earl DA, vonHoldt BM. 2012. STRUCTURE HARVESTER: A website and program for visualizing STRUCTURE output and implementing the Evanno method. Conservation Genetics Resources 4:359–361. DOI: 10.1007/s12686-011-9548-7.

Estes JA, Terborgh J, Brashares JS, Power ME, Berger J, Bond WJ, Carpenter SR, Essington TE, Holt RD, Jackson JBC, Marquis RJ, Oksanen L, Oksanen T, Paine RT, Pikitch EK, Ripple WJ, Sandin SA, Scheffer M, Schoener TW, Shurin JB, Sinclair ARE, Soulé ME, Virtanen R, Wardle DA. 2011. Trophic Downgrading of Planet Earth. Science 333:301–306. DOI: 10.1126/science.1205106.

Excoffier L, Laval G, Schneider S. 2005. Arlequin ver 3.1: An Integrated Software Package for Population Genetics Data Analysis. DOI: 10.1093/nq/s9-II.43.326-b.

Falush D, Stephens M, Prithard JK. 2003. Inference of Population Structure Using Multilocus Genotype Data: Linked Loci and Correlated Allele Frequencies Daniel. JAMA: The Journal of the American Medical Association 164:1567–1587. DOI: 10.1001/jama.1987.03400040069013.

Garza JC, Williamson EG. 2001. Detection of reduction in population size using data from microsatellite loci. Molecular Ecology 10:305–318.

Gavashelishvili A, Lukarevskiy V. 2008. Modelling the habitat requirements of leopard Panthera pardus in west and central Asia. Journal of Applied Ecology 45:579–588. DOI: 10.1111/j.1365-2664.2007.01432.x.

Goossens B, Chikhi L, Ancrenaz M, Lackman-Ancrenaz I, Andau P, Bruford MW. 2006. Genetic Signature of Anthropogenic Population Collapse in Orang-utans. PLoS Biology 4:e25. DOI: 10.1371/journal.pbio.0040025.

Grant T. 2012. Leopard population density, home range size and movement patterns in a mixed landuse area of the mangwe district of Zimbabwe.: 1–143.

Hájková P, Pertoldi C, Zemanová B, Roche K, Hájek B, Bryja J, Zima J. 2007. Genetic structure and evidence for recent population decline in Eurasian otter populations in the Czech and Slovak Republics: implications for conservation. Journal of Zoology 272:1–9. DOI: 10.1111/j.1469-7998.2006.00259.x.

Harihar A, Pandav B, Goyal SP. 2009. Responses of tiger (Panthera tigris) and their prey to removal of anthropogenic influences in Rajaji National Park, India. European Journal of Wildlife Research 55:97–105. DOI: 10.1007/s10344-008-0219-2.

Harihar A, Pandav B, Goyal SP. 2011. Responses of leopard Panthera pardus to the recovery of a tiger Panthera tigris population. Journal of Applied Ecology 48:806–814. DOI: 10.1111/j.1365-2664.2011.01981.x.

Hines, J. E. (2006). Program presence. See http://www.mbrpwrc.usgs.gov/software/doc/presence/presence.html.

Hu Y, Guo Y, Qi D, Zhan X, Wu H, Bruford MW, Wei F. 2011. Genetic structuring and recent demographic history of red pandas (Ailurus fulgens) inferred from microsatellite and mitochondrial DNA. Molecular Ecology 20:2662–2675. DOI: 10.1111/j.1365-294X.2011.05126.x.

Jacobson AP, Gerngross P, Lemeris Jr. JR, Schoonover RF, Anco C, Breitenmoser-Würsten C, Durant SM, Farhadinia MS, Henschel P, Kamler JF, Laguardia A, Rostro-García S, Stein AB, Dollar L. 2016. Leopard (Panthera pardus) status, distribution, and the research efforts across its range. PeerJ 4:e1974. DOI: 10.7717/peerj.1974.

Jhala YV, Qureshi Q, Gopal R. 2015. The status of tigers in India, 2014.

Johnsingh AJT, Ramesh K, Qureshi Q, A. D, Goyal SP, Rawat GS, Rajapandian K, Prasad S. 2004. Conservation Status of Tiger and Associated Species in the Terai Arc Landscape, India.

Kalinowski ST, Taper ML, Marshall TC. 2007. Revising how the computer program cervus accommodates genotyping error increases success in paternity assignment. Molecular Ecology 16:1099–1106. DOI: 10.1111/j.1365-294X.2007.03089.x.

Kalle R, Ramesh T, Qureshi Q, Sankar K. 2011. Density of tiger and leopard in a tropical deciduous forest of Mudumalai Tiger Reserve, southern India, as estimated using photographic capture–recapture sampling. Acta Theriologica 56:335–342. DOI: 10.1007/s13364-011-0038-9.

Karanth KU, Chellam R. 2009. Carnivore conservation at the crossroads. Oryx 43:1. DOI: 10.1017/S003060530843106X.

Karanth KK, Gopalaswamy AM, DeFries R, Ballal N. 2012. Assessing Patterns of Human-Wildlife Conflicts and Compensation around a Central Indian Protected Area. PLoS ONE 7:e50433. DOI: 10.1371/journal.pone.0050433.

Karanth KK, Naughton-Treves L, DeFries R, Gopalaswamy AM. 2013. Living with Wildlife and Mitigating Conflicts Around Three Indian Protected Areas. Environmental Management 52:1320–1332. DOI: 10.1007/s00267-013-0162-1.

Karanth KK, Nichols JD, Hines JE, Karanth KU, Christensen NL. 2009. Patterns and determinants of mammal species occurrence in India. Journal of Applied Ecology 46:1189–1200. DOI: 10.1111/j.1365-2664.2009.01710.x.

Karanth KK, Nichols JD, Karanth KU, Hines JE, Christensen NL. 2010. The shrinking ark: patterns of large mammal extinctions in India. Proceedings of the Royal Society B: Biological Sciences 277:1971–1979. DOI: 10.1098/rspb.2010.0171.

Karanth KU, Sunquist M. 2000. Behavioural Correlates of Predation by Tiger (Panthera tigris) & Leopard (Panthera pardus) in. The zoological society of London 4:255–265.

Lorenzen ED, Nogués-Bravo D, Orlando L, Weinstock J, Binladen J, Marske KA, Ugan A, Borregaard MK, Gilbert MTP, Nielsen R, Ho SYW, Goebel T, Graf KE, Byers D, Stenderup JT, Rasmussen M, Campos PF, Leonard JA, Koepfli K-P, Froese D, Zazula G, Stafford TW, Aaris-Sørensen K, Batra P, Haywood AM, Singarayer JS, Valdes PJ, Boeskorov G, Burns JA, Davydov SP, Haile J, Jenkins DL, Kosintsev P, Kuznetsova T, Lai X, Martin LD, McDonald HG, Mol D, Meldgaard M, Munch K, Stephan E, Sablin M, Sommer RS, Sipko T, Scott E, Suchard MA, Tikhonov A, Willerslev R, Wayne RK, Cooper A, Hofreiter M, Sher A, Shapiro B, Rahbek C, Willerslev E. 2011. Species-specific responses of Late Quaternary megafauna to climate and humans. Nature 479:359–364. DOI: 10.1038/nature10574.

Marie J, Luikart G. 1996. Power Analysis of Two Tests for Detecting Recent Population Bottlenecks From Allele Frequency Data. Test 144:2001–2014. DOI: Article.

Maroju PA, Yadav S, Kolipakam V, Singh S, Qureshi Q, Jhala Y. 2016. Schrodinger’s scat: a critical review of the currently available tiger (Panthera Tigris) and leopard (Panthera pardus) specific primers in India, and a novel leopard specific primer. BMC Genetics 17:37. DOI: 10.1186/s12863-016-0344-y.

Marsden CD, Woodroffe R, Mills MGL, Mcnutt JW, Creel S, Groom R, Emmanuel M, Cleaveland S, Kat P, Rasmussen GSA, Ginsberg J, Lines R, Andre J-M, Begg C, Wayne RK, Mable BK. 2012. Spatial and temporal patterns of neutral and adaptive genetic variation in the endangered African wild dog (*Lycaon pictus*). Molecular Ecology 21:1379–1393. DOI: 10.1111/j.1365-294X.2012.05477.x.

Martinez-Cruz B, Godoy JA, Negro JJ. 2007. Population fragmentation leads to spatial and temporal genetic structure in the endangered Spanish imperial eagle. Molecular Ecology 16:477–486. DOI: 10.1111/j.1365-294X.2007.03147.x.

Miquel C, Bellemain E, Poillot C, Bessière J, Durand A, Taberlet P. 2006. Quality indexes to assess the reliability of genotypes in studies using noninvasive sampling and multiple-tube approach. Molecular Ecology Notes 6:985–988. DOI: 10.1111/j.1471-8286.2006.01413.x.

Mondal K, Sandar K, Qureshi Q, Gupta S, Chourasia P. 2012. Estimation of population and survivorship of leopard through photographic cature-recapture sampling in west india. World Journal of Zoology 7:30–39. DOI: 10.5829/idosi.wjz.2012.7.

Mondol S, Bruford MW, Ramakrishnan U. 2013. Demographic loss, genetic structure and the conservation implications for Indian tigers. Proceedings of the Royal Society B: Biological Sciences 280:20130496–20130496. DOI: 10.1098/rspb.2013.0496.

Mondol S, Karanth KU, Ramakrishnan U. 2009. Why the Indian Subcontinent Holds the Key to Global Tiger Recovery. PLoS Genetics 5:e1000585. DOI: 10.1371/journal.pgen.1000585.

Mondol S, Kumar NS, Gopalaswamy A, Sunagar K, Karanth KU, Ramakrishnan U. 2014. Identifying species, sex and individual tigers and leopards in the Malenad-Mysore Tiger Landscape, Western Ghats, India. Conservation Genetics Resources 7:353–361. DOI: 10.1007/s12686-014-0371-9.

Mondol S, R N, Athreya V, Sunagar K, Selvaraj VM, Ramakrishnan U. 2009. A panel of microsatellites to individually identify leopards and its application to leopard monitoring in human dominated landscapes. BMC Genetics 10:79. DOI: 10.1186/1471-2156-10-79.

Mondol S, Sridhar V, Yadav P, Gubbi S, Ramakrishnan U. 2015. Tracing the geographic origin of traded leopard body parts in the indian subcontinent with DNA-based assignment tests. Conservation Biology 29:556–564. DOI: 10.1111/cobi.12393.

Mora MS, Mapelli FJ, Gaggiotti OE, Kittlein MJ, Lessa EP. 2010. Dispersal and population structure at different spatial scales in the subterranean rodent Ctenomys australis. BMC Genetics 11:9. DOI: 10.1186/1471-2156-11-9.

Naha D, Sathyakumar S, Rawat GS. 2018. Understanding drivers of human-leopard conflicts in the Indian Himalayan region: Spatio-temporal patterns of conflicts and perception of local communities towards conserving large carnivores. PLOS ONE 13:e0204528. DOI: 10.1371/journal.pone.0204528.

Nater A, Greminger MP, Arora N, van Schaik CP, Goossens B, Singleton I, Verschoor EJ, Warren KS, Krützen M. 2015. Reconstructing the demographic history of orang-utans using Approximate Bayesian Computation. Molecular Ecology 24:310–327. DOI: 10.1111/mec.13027.

Van Oosterhout C, Hutchinson WF, Wills DPM, Shipley P. 2004. MICRO-CHECKER: software for identifying and correcting genotyping errors in microsatellite data. Molecular Ecology Notes 4:535–538. DOI: 10.1111/j.1471-8286.2004.00684.x.

Peakall R, Smouse PE. 2012. GenAlEx 6.5: genetic analysis in Excel. Population genetic software for teaching and research--an update. Bioinformatics 28:2537–2539. DOI: 10.1093/bioinformatics/bts460.

Perez I, Geffen E, Mokady O. 2006. Critically Endangered Arabian leopards Panthera pardus nimr in Israel: estimating population parameters using molecular scatology. Oryx 40:295–301. DOI: 10.1017/S0030605306000846.

Piry S, Alapetite A, Cornuet JM, Paetkau D, Baudouin L, Estoup A. 2004. GENECLASS2: A software for genetic assignment and first-generation migrant detection. Journal of Heredity 95:536–539. DOI: 10.1093/jhered/esh074.

Prithard JK, Stephes M, Donnelly P. 2000. Inference of population structure using multilocus genotype data. Genetics Society of America 155:945–959. DOI: 10.1111/j.1471-8286.2007.01758.x.

Rangarajan, M. (2005). India’s wildlife history: an introduction. Orient Blackswan.

Raza RH, Chauhan DS, Pasha MKS, Sinha S. 2012. Illuminating the Blind Spot: A study on illegal trade in Leopard parts in India (2001-2010).

Ripple WJ, Estes JA, Beschta RL, Wilmers CC, Ritchie EG, Hebblewhite M, Berger J, Elmhagen B, Letnic M, Nelson MP, Schmitz OJ, Smith DW, Wallach AD, Wirsing AJ. 2014. Status and Ecological Effects of the World’s Largest Carnivores. Science 343:1241484. DOI: 10.1126/science.1241484.

Roy PS, Joshi PK, Singh S, Agarwal S, Yadav D, Jegannathan C. 2006. Biome mapping in India using vegetation type map derived using temporal satellite data and environmental parameters. Ecological Modelling 197:148–158. DOI: 10.1016/j.ecolmodel.2006.02.045.

Seidensticker, J. 1990. Leopards living at the edge of the Royal Chitwan National Park, Nepal. Conservation in developing countries: problems and prospects:415–423.

Selvan KM, Lyngdoh S, Habib B, Gopi GV. 2014. Population density and abundance of sympatric large carnivores in the lowland tropical evergreen forest of Indian Eastern Himalayas. Mammalian Biology 79:254–258. DOI: 10.1016/j.mambio.2014.03.002.

Sergio F, Caro T, Brown D, Clucas B, Hunter J, Ketchum J, McHugh K, Hiraldo F. 2008. Top Predators as Conservation Tools: Ecological Rationale, Assumptions, and Efficacy. Annual Review of Ecology, Evolution, and Systematics 39:1–19. DOI: 10.1146/annurev.ecolsys.39.110707.173545.

Sillero-Subiri C, Laurenson K. 2001. Interactions between carnivores and local communities: Conflict of co-existence? Carnivore conservation:282–312.

Spong G, Johansson M, Bjorklund M. 2000. High genetic variation in leopards indicates large and long-term stable effective population size. Molecular Ecology 9:1773–1782. DOI: 10.1046/j.1365-294X.2000.01067.x.

Stein, A. B., Athreya, V., Gerngross, P., Balme, G., Henschel, P., Karanth, U., … & Laguardia, A. (2016). Panthera pardus, Leopard. The IUCN Red List of Threatened Species 2016: e.T15954A50659089.

Storz JF, Beaumont MA. 2002. Testing for Genetic Evidence of Population Expansion and Contraction: An Empirical Analysis of Microsatellite DNA Variation Using a Hierarchical Bayesian Model Published by: Society for the Study of Evolution Stable URL: http://www.jstor.org/stable/306. Evolution 56:154–166.

Sugimoto T, Gray TNE, Higashi S, Prum S. 2014. Examining genetic diversity and identifying polymorphic microsatellite markers for noninvasive genetic sampling of the Indochinese leopard (Panthera pardus delacouri). Mammalian Biology 79:406–408. DOI: 10.1016/j.mambio.2014.06.002.

Sunquist ME. 1983. Dispersal of Three Radiotagged Leopards. Journal of Mammalogy 64:337–341.

Terborgh J, Lopez L, V PN, Rao M, Orihuela G, Riveros M, Ascanio R, Adler GH, Thomas D, Balbas L, Terborgh J, Lopez L, V PN, Rao M, Shahabuddin G, Orihuela G, Riveros M, Ascanio R, Adler GH. 2001. Ecological Meltdown in Predator-Free Forest Fragments Published by: American Association for the Advancement of Science Linked references are available on JSTOR for this article: Ecological Meltdown in Predator-Free Forest Fragments. Science (New York, N.Y.) 294:1923–1926. DOI: 10.1126/science.1064397.

Thapa K, Shrestha R, Karki J, Thapa GJ, Subedi N, Pradhan NMB, Dhakal M, Khanal P, Kelly MJ. 2014. Leopard Panthera pardus fusca Density in the Seasonally Dry, Subtropical Forest in the Bhabhar of Terai Arc, Nepal. Advances in Ecology 2014:1–12. DOI: 10.1155/2014/286949.

Tucker JM, Schwartz MK, Truex RL, Pilgrim KL, Allendorf FW. 2012. Historical and Contemporary DNA Indicate Fisher Decline and Isolation Occurred Prior to the European Settlement of California. PLoS ONE 7:e52803. DOI: 10.1371/journal.pone.0052803.

Uphyrkina O, Johnson WE, Quigley H, Miquelle D, Marker L, Bush M, O’Brien SJ. 2001. Phylogenetics, genome diversity and origin of modern leopard, Panthera pardus\ndoi:10.1046/j.0962-1083.2001.01350.x. Molecular Ecology 10:2617–2633.

Valdiosera CE, García-Garitagoitia JL, Garcia N, Doadrio I, Thomas MG, Hänni C, Arsuaga J-L, Barnes I, Hofreiter M, Orlando L, Götherström A. 2008. Surprising migration and population size dynamics in ancient Iberian brown bears (Ursus arctos). Proceedings of the National Academy of Sciences 105:5123–5128. DOI: 10.1073/pnas.0712223105.

Valiere N. 2002. Gimlet: a computer program for analysing genetic individual identification data. Molecular Ecology Notes 2:377–379. DOI: 10.1046/j.1471-8286.2002.00228.x.

Wang SW, Macdonald DW. 2009. The use of camera traps for estimating tiger and leopard populations in the high altitude mountains of Bhutan. Biological Conservation 142:606–613. DOI: 10.1016/j.biocon.2008.11.023.

Wolf C, Ripple WJ. 2017. Range contractions of the world’s large carnivores. Royal Society Open Science 4:170052. DOI: 10.1098/rsos.170052.

Yang J, Jiang Z. 2011. Genetic diversity, population genetic structure and demographic history of Przewalski’s gazelle (*Procapra przewalskii*): Implications for conservation. Conservation Genetics 12:1457–1468. DOI: 10.1007/s10592-011-0244-7.

Zhu L, Zhan X, Wu H, Zhang S, Meng T, Bruford MW, Wei F. 2010. Conservation Implications of Drastic Reductions in the Smallest and Most Isolated Populations of Giant Pandas. Conservation Biology 24:1299–1306. DOI: 10.1111/j.1523-1739.2010.01499.x.

