## Supplementary Figure and Table for "Genetic analyses reveal population structure and recent decline in leopards (*Panthera pardus fusca*) across Indian subcontinent"

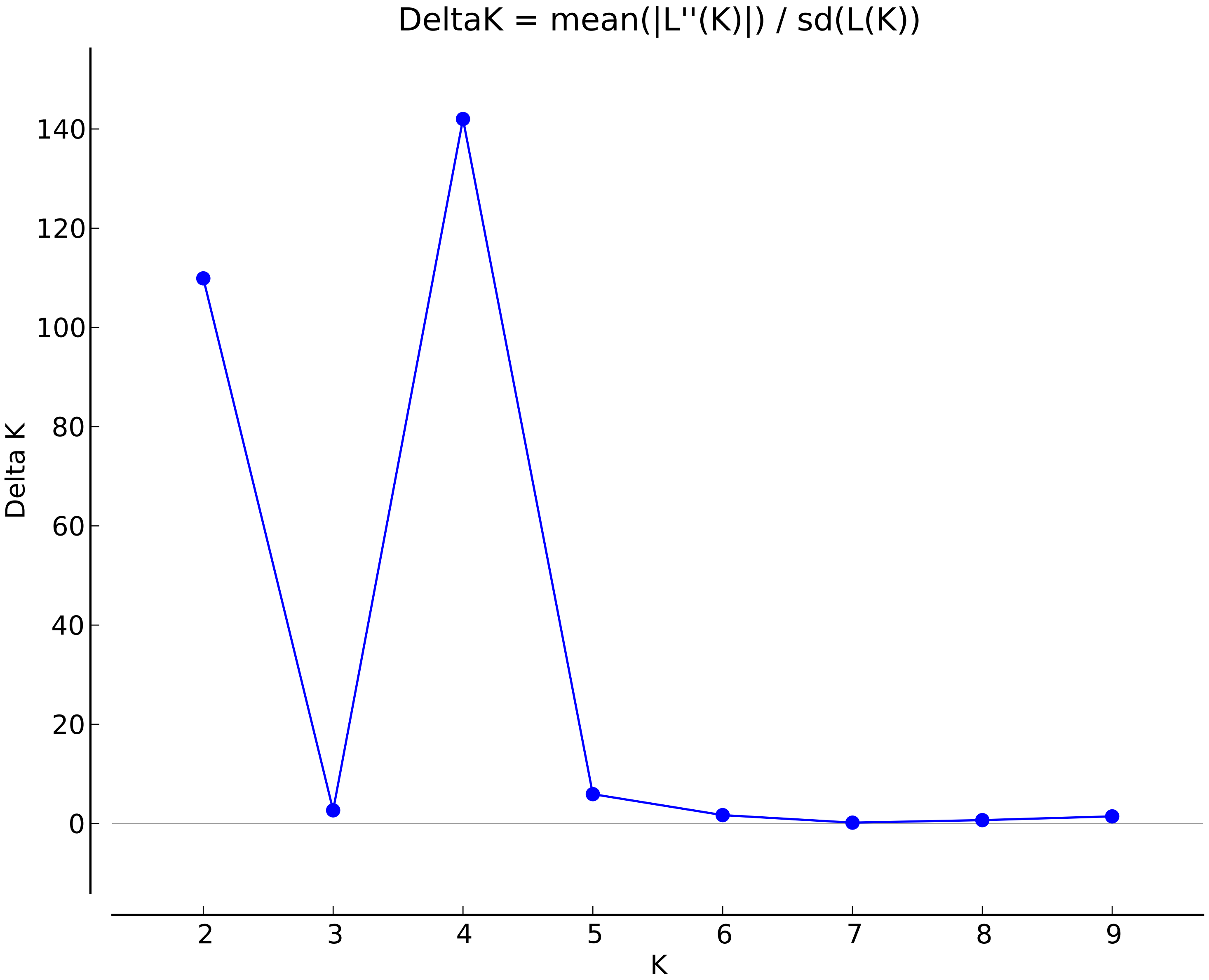
DeltaK = mean(|L''(K)|) / sd(L(K))
Delta K
K
0
20
40
60
80
100
120
140
2
3
4
5
6
7
8
9

Supplementary Table 1– Prior models used for leopard demography analyses in Storz and Beaumont method. A generation time of 5 years is used.

| **Runs** | **log($N_0$)** | **log($N_1$)** | **log(\)** | **log($T$)** | **log($N_0$)** | **log($N_1$)** | **log(\)** | **log($T$)** |
|---|---|---|---|---|---|---|---|---|
| **Run 01** | **5 1** | **5 1** | **-3.5 1** | **3 1** | **3.5 2 0 0.5** | **6 3 0 0.5** | **-3.5 0.25 0 0.5** | **3 2 0 0.5** |
| **Run 02** | **5 1** | **5 1** | **-3.5 1** | **3 1** | **3.5 2 0 0.5** | **5 2 0 0.5** | **-3.5 0.25 0 0.5** | **3 2 0 0.5** |
| **Run 03** | **5 1** | **5 1** | **-3.5 1** | **3 1** | **3.5 2 0 0.5** | **5 1 0 0.5** | **-3.5 0.25 0 0.5** | **3 2 0 0.5** |
| **Run 04** | **5 1** | **5 1** | **-3.5 1** | **3 1** | **3.5 2 0 0.5** | **6 1 0 0.5** | **-3.5 0.25 0 0.5** | **3.5 2 0 0.5** |
| **Run 05** | **5 1** | **5 1** | **-3.5 1** | **3 1** | **4 2 0 0.5** | **6 2 0 0.5** | **-3.5 0.25 0 0.5** | **3 2 0 0.5** |
| **Run 06** | **5 1** | **5 1** | **-3.5 1** | **3 1** | **3.5 2 0 0.5** | **5 1 0 0.5** | **-3.5 0.25 0 0.5** | **3 2 0 0.5** |
